# Nanoparticle-induced augmentation of neutrophils’ phagocytosis of bacteria

**DOI:** 10.1101/2022.05.14.491866

**Authors:** Kathryn M. Rubey, Alexander R. Mukhitov, Jia Nong, Jichuan Wu, Vera P. Krymskaya, Jacob W. Myerson, G. Scott Worthen, Jacob S. Brenner

## Abstract

Despite the power of antibiotics, bacterial infections remain a major killer, due to antibiotic resistance and hosts with dysregulated immune systems. We and others have been developing drug-loaded nanoparticles that home to the sites of infection and inflammation via engineered tropism for neutrophils, the first-responder leukocytes in bacterial infections. Here, we examined how a member of a broad class of neutrophil-tropic nanoparticles affects neutrophil behavior, specifically questioning whether the nanoparticles attenuate an important function, bacterial phagocytosis. We found these nanoparticles actually *augment* phagocytosis of non-opsonized bacteria, increasing it by ~50%. We showed this augmentation of phagocytosis is likely co-opting an evolved response, as opsonized bacteria also augment phagocytosis of non-opsonized bacteria. Enhancing phagocytosis of non-opsonized bacteria may prove particularly beneficial in two clinical situations: in hypocomplementemic patients (meaning low levels of the main bacterial opsonins, complement proteins, seen in conditions such as neonatal sepsis and liver failure) or for bacteria that are largely resistant to complement opsonization (e.g., *Neisseria*). Additionally, we observe that; a) prior treatment with bacteria augments neutrophil uptake of neutrophil-tropic nanoparticles; b) neutrophil-tropic nanoparticles colocalize with bacteria inside of neutrophils. The observation that neutrophil-tropic nanoparticles enhance neutrophil phagocytosis and localize with bacteria inside neutrophils suggests that these nanoparticles will serve as useful carriers for drugs to ameliorate bacterial diseases.

## 1 Introduction

The introduction of antibiotics last century has left the lay public thinking that bacterial infections are a relatively solved problem, but the clinical reality is that these diverse diseases still cause a huge number of deaths and severe illnesses, even when antibiotics are used (*Pneumonia,* no date; File and Marrie, 2010; Xu *et al.*, 2020; Rubey and Brenner, 2021). The reasons for incomplete effectiveness include at least 3 major factors: First, the “host response” to bacteria is often deleterious, epitomized by sepsis-induced organ failure(Pechous, 2017; Lelubre and Vincent, 2018). Second, bacterial resistance to antibiotics is rising precipitously(Boucher *et al.*, 2009; Reardon, 2015; Centers for Disease Control and, 2019). Third, many hosts, by virtue of age, underlying condition, or therapeutic regimen, are immunocompromised and unable to clear bacteria even in the presence of antibiotics(Grant and Hung, 2013; Tosi *et al.*, 2018). All of these problems are compounded by a lack of innovation in therapies compared to other fields(Silver, 2011).

To solve this problem, we and others have been developing nanoparticles that can deliver drugs directly to the site of infection (Rubey and Brenner, 2021). This approach may address each of the 3 problems listed above, depending on whether the cargo drug is an antibiotic, whose therapeutic index can be improved by localization, or an immunomodulator. A particularly promising method for targeting these nanoparticles to acute infections is directing the nanoparticles to neutrophils. Neutrophils are “first responder” leukocytes for most acute bacterial infections, and massively accumulate in sites of infection(Yipp *et al.*, 2017). By delivering antibiotics to neutrophils, nanoparticles could improve bacterial killing. This may be especially useful for antibiotics that cannot cross neutrophil membranes, such as aminoglycosides. Alternatively, neutrophil-tropic nanoparticles could deliver anti-inflammatories to modulate some of the neutrophils’ more deleterious responses (part of the dysregulated host defense of sepsis (Zemans, Colgan and Downey, 2009)), such as production of tissue damaging mediators such as neutrophil extracellular traps (NETs)(Yipp and Kubes, 2013), reactive oxygen species (ROS), and proteases (Moraes, Zurawska and Downey, 2006). Nanoparticles with tropism for neutrophils have potential to greatly improve most key aspects of acute bacterial infections, and even non-bacterial inflammatory diseases, such as acute respiratory distress syndrome (ARDS), the highly neutrophilic lung inflammation that kills in COVID-19(Sinha *et al.,* 2022).

We recently reported a nanomaterials screen to identify nanoparticles with strong tropism to the neutrophils that accumulate in the capillaries of the lungs during inflammation and play a major role in pneumonia, COVID-19, and ARDS(Myerson *et al.*, 2022). We found a broad class of nanoparticles with neutrophil-tropism: “nanoparticles with agglutinated proteins” (NAPs). NAPs have surface-accessible proteins arranged in a non-crystalline pattern (meaning agglutinated / amorphous). By contrast, crystalline protein nanoparticles (viral capsids, ferritin, etc), which have their surface proteins arranged in a fixed and regular pattern, do not have neutrophil tropism. We showed this tropism is due to the fact that NAPs rapidly bind the serum proteins, including C3b and other complement proteins, suggesting complement activation, while non-NAPs do not. Complement is an important component of the immune system that aids phagocytic cells in recognizing particulate matter to be phagocytosed. Previous studies have been dedicated to further understanding of complement binding and activation caused by nanoparticles as it can be a barrier to a nanocarrier’s therapeutic potential (Scieszka *et al.*, 1991; Inturi *et al.*, 2015; Chen *et al.*, 2017; Moghimi and Simberg, 2017; Betker *et al.*, 2018; Vu *et al.*, 2019). We have chosen to use this property to our advantage as serum-opsonization of NAPs is essential for their strong neutrophil tropism.

To develop NAPs for targeted delivery to neutrophils in infectious diseases, such as pneumonia, we aim here to ensure that NAPs coordinate with the key beneficial functions of neutrophils, and do not negatively impact their function. Previous studies have shown that nanoparticle phagocytosis may decrease neutrophil adhesion and migration(Fromen *et al.*, 2017). Probably the most essential function of neutrophils during infections like pneumonia is phagocytosis of bacteria, since phagocytosis is necessary for killing of certain bacteria(Lee, Harrison and Grinstein, 2003). Here, we tested neutrophil phagocytosis of NAPs before and after neutrophil phagocytosis of the common bacterial pathogen, *E. coli.* We found that NAPs do not negatively impact neutrophil phagocytosis of bacteria and NAPs localize to neutrophils that have also taken up bacteria. Quite surprisingly, NAPs, given *after* bacteria, enhance efficiency of neutrophils’ phagocytosis of bacteria that have not been opsonized by serum proteins. This effect, termed here *second particle augmentation factor* (*2PAF*) was illustrated in both: flow cytometry and microscopy. Our results suggest applicability of NAPs to two important clinical situations: hypocomplementemic states (e.g., the neonate (Wolach *et al.*, 1994; Kemp and Campbell, 1996; Schelonka and Infante, 1998; McGreal, Hearne and Spiller, 2012; Zimmermann and Jones, 2021) and liver failure) and bacteria that have strong complement-defense mechanisms(Flannagan, Cosío and Grinstein, 2009) (e.g., *Neisseria* coats itself in the complement inhibitor Factor H). In such clinical situations, NAPs’ augmentation of bacterial phagocytosis and colocalization with bacteria in neutrophils could provide a major benefit, beyond the benefits of cargo drugs themselves.

## 2 Results and Discussion

We recently developed a diverse class of nanoparticles with neutrophil-tropism: NAPs(Myerson *et al.*, 2022). In the present study, we focus on a prototypical member of this class, lysozyme-dextran nanogels (hereafter referred to as “nanogels” or NGs). NGs have the benefit for antibiotic delivery of prolonged nanoparticle shelf-life (years at 4C) and a very high drug-to-carrier mass ratio(Myerson *et al.*, 2018, 2019).

In these experiments with nanogels, we again confirmed our previous findings that particle uptake is enhanced by opsonization by complement proteins present in serum(Myerson *et al.*, 2022). When complement protein C3 is depleted from serum via cobra venom factor (CVF)(Haihua *et al.*, 2018), we see a significant decrease in the percent of neutrophils that take up nanogels (Supplemental Figure 1). The green serum nanogels (second green bar) showed ~80% positivity, as compared to the green CVF nanogels (third and fifth green bars) which had ~30% positivity. It has been well established that complement binding to *E. coli* is necessary for neutrophil phagocytosis of the bacteria(Horwitz and Silverstein, 1980; Brekke *et al.*, 2007). We utilized serum-opsonization of both NAPs and *E. coli* bioparticles for these experiments.

One of the first tests of whether these nanocarriers can be used to augment bacterial killing is whether neutrophils will take up the nanocarriers after having been exposed to bacteria. In this experiment, diagrammed in Figure 1A, neutrophils were incubated with heat-killed *E. coli* bioparticles which have surface conjugated pHrodo green, a pH-sensitive dye that fluoresces green only when in the low pH environment of the phagosome. After this 60-minute 37°C incubation, the neutrophils were pelleted and washed to remove free bacteria. The neutrophils were then incubated with nanogels for 15 minutes, and subjected to flow cytometry. Before exposure to neutrophils, half the samples of *E. coli* were serum-opsonized (hereafter referred to as “Serum EC”), while the other half were not exposed to serum (simply labeled as “EC” in Fig 1). Similarly, nanogels were divided into serum-opsonized (“Serum NG”) and not (simply “NG”).

**Figure 1.**
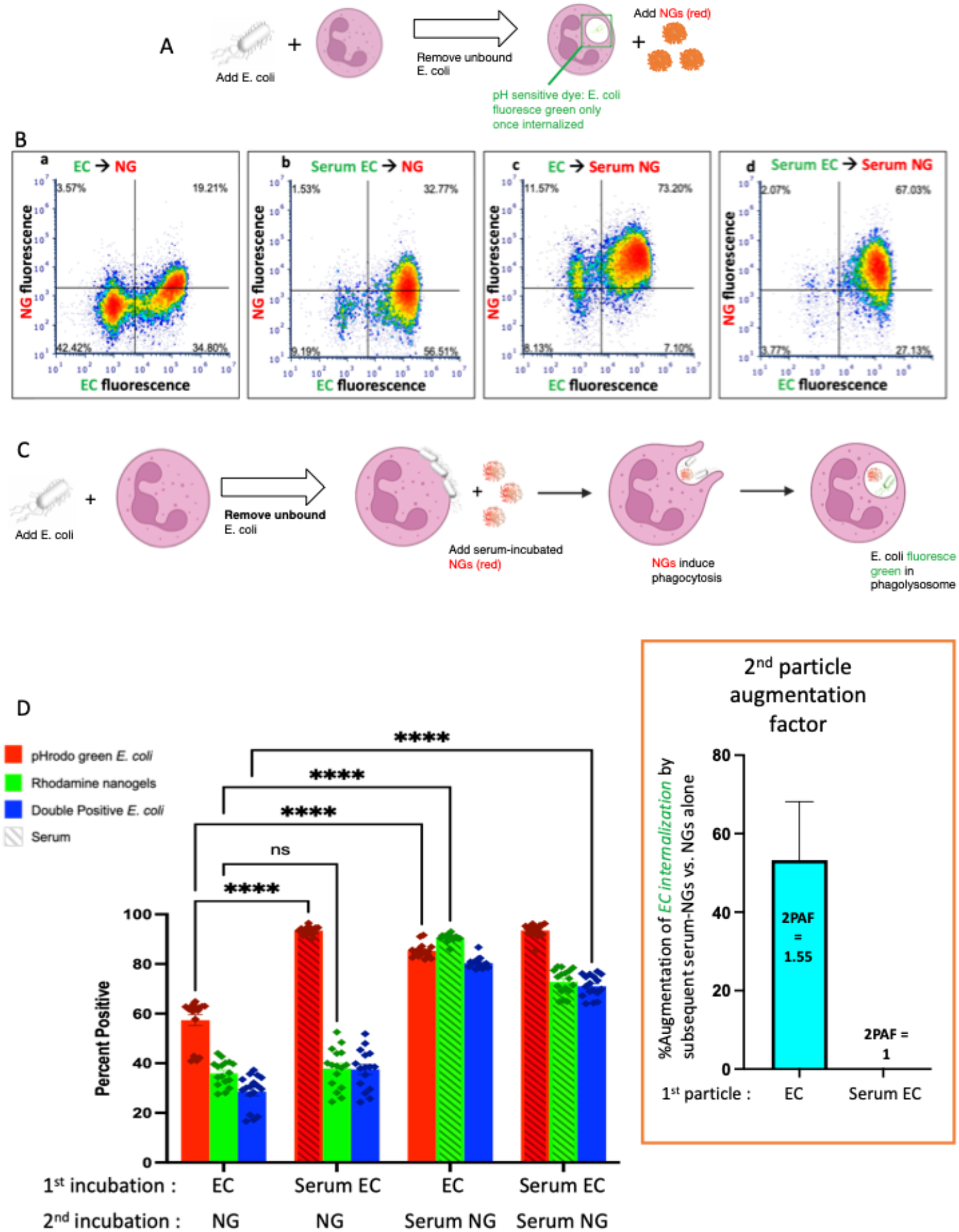
Nanogels augment phagocytosis of *E. coli.* A) Neutrophils were incubated with pH-sensitive fluorescent *E. coli* that only fluoresce in the low pH of the phagosome, thus allowing detection of their phagocytosis. Notably, *E. coli* that bind to the exterior of neutrophils but are not internalized do *not* fluoresce. First neutrophils were exposed to green *E. coli,* thoroughly washed to remove unbound *E. coli,* and then incubated with red nanogels (“NG”). Importantly, before exposure to the neutrophils, select aliquots of nanogels and *E. coli* were opsonized by serum and thoroughly washed before exposure to neutrophils. B) Representative dot-plots of the neutrophils subjected to flow cytometry *after* the second incubation, examining for green fluorescence (x-axis, indicating phagocytosis of the *E. coli*) and red (y-axis, nanogels). At the top of each subpanel, we indicate whether the *E. coli* (EC) were not exposed to serum (simply “EC”) or were serum-opsonized (“Serum EC”); similar notation is used for nanogels without serum exposure (simply “NG”) or those that were serum-opsonized (“Serum NG”). Neutrophils positive for uptake of green *E. coli* are represented as dots to the right of the vertical bar in the middle of the plot, with that threshold determined by measuring fluorescence of *E. coli* not exposed to any *E. coli.* Similarly, neutrophils represented as dots above the horizontal line are positive for red nanogels C) Diagram of the proposed mechanism underlying this 2PFE of 1.5. Some *E. coli* remain bound to the neutrophil surface even after thoroughly washing the cells (second neutrophil image), but these surface-bound *E. coli* do not fluoresce since buffer pH of ~7 prevents their pH-sensitive dye from fluorescing. When these neutrophils encounter Serum NG, they phagocytose not just the nanogels, but also the *E. coli* (third neutrophil image), causing the *E. coli* to fluoresce in the low pH of the phagosome (fourth neutrophil). D) Quantification of the flow cytometry experiments with n=6 biological replicates for each sample. The bottom axis lists for the 1st and 2nd incubations whether the *E. coli* and nanogels were first exposed to serum or not. The y-axis lists the % of neutrophils that were positive for phagocytosed *E. coli* (recalling that pH-sensitive E.coli on the *surface* of neutrophils will *not* fluoresce). Among the many bars to compare, the most important is comparing the first vs third red bars, which both measure the % of neutrophils that phagocytosed non-serum-opsonized *E. coli* (EC). When the second incubation was with NG (not serum exposed), only 55% of neutrophils were positive for the EC presented during the first incubation. However, when the second incubation was with *Serum* NG, 85% of neutrophils were positive for the EC presented during the first incubation. This important ratio (first vs third red bar), which we call the “2nd particle augmentation factor (2PAF)” was ~1.55 (*inset in box,* red), and *measures the fold-increase in phagocytosis of ECs induced by Serum NG (compared to non-serum-exposed NG, which serves as an internal control*). Notably, while Serum NG augment uptake of (non-opsonized) EC by 50% (2PAF of 1.5), Serum NG do not significantly augment neutrophil uptake of Serum EC (second vs fourth red bar, 2PAF = 1.0, blue bar in inset). Thus, Serum NG only augment phagocytosis of non-opsonized *E. coli.*

Flow cytometry was gated to analyze neutrophils exclusively (Supplemental Figure 2). Representative flow cytometry dot-plots are depicted in Figure 1B and quantified in n=6 biological replicates in Figure 1D. We compared the summary statistic of the percentage of neutrophils positive for *E. coli* and/or nanogel fluorescence. Among the numerous comparisons that can be made in this dataset, a surprising finding is demonstrated by comparing the first and third *E. coli* uptake values (red bars) in Figure 1D. These conditions measure the fraction of neutrophils that are positive for phagocytosis of *E. coli* that had not been exposed to serum (“EC”). When these neutrophils were incubated with nanogels that had not been exposed to serum (first red bar), 55% of the neutrophils were positive for *E. coli* phagocytosis. This percentage of neutrophils positive for *E. coli* went up to 85% if the nanogels had been pre-opsonized by serum. This means that serum-opsonized nanogels are able to augment *E. coli* phagocytosis. This augmentation occurred even though the nanogels were delivered *after* free *E. coli* had been washed away from the neutrophils.

We term this enhancement the “second particle augmentation factor” (2PAF) and define it for this particular experiment (Fig 1) as the following ratio: (% neutrophils positive for EC phagocytosis when the second delivered particle is Serum NG) / (% neutrophils positive for EC phagocytosis when the second delivered particle is [non-serum-exposed] NG). 2PAF can be more generally defined as: (% neutrophils positive for particle #1 phagocytosis, given that particle #2 is serum-opsonized) / (% neutrophils positive for particle #1 phagocytosis, given that particle #2 was not exposed to serum); where particle #1 refers to the particle (or microbe) neutrophils are exposed to in the first incubation, and particle #2 refers to the second incubation. Thus, a 2PAF > 1 indicates that serum-opsonized particle #2 are able to augment phagocytosis of particle #1 (as compared to the control condition, which uses particle #2 that was not exposed to serum). Calculating the 2PAF for non-serum exposed *E. coli* (EC), we thus get 2PAF = 85% / 55% = 1.55, meaning that serum-opsonized nanogels increase neutrophil phagocytosis of these *E. coli* by 50% (Fig 1D, inset, blue bar). A 2PAF > 1 is only found when the *E. coli* have not been serum opsonized: 2PAF = 1 when using serum-opsonized *E. coli,* a 0% increase (Fig 1D, inset). Thus, serum-opsonized nanogels are able to augment uptake of non-opsonized bacteria, but not opsonized bacteria, the latter of which are already phagocytosed so extensively (~100%) that we cannot detect improvement within the dynamic range of this assay. The mechanism by which serum enhances NAP phagocytosis is through coating the particles with complement proteins (e.g., C3), as shown previously (Myerson *et al.*, 2022), and in Supplementary Figure 1. This suggests that serum-opsonized nanogels would be most effective in augmenting phagocytosis of bacteria that are not complement-opsonized, such as bacteria that evade complement by covering themselves with Factor H (*Neisseria*, etc), or bacterial infections in hypocomplementemic hosts (e.g., neonatal sepsis).

A hypothesis to explain this enhancement is outlined in Figure 1C. The neutrophils are first incubated with green *E. coli* that fluoresce only when in the phagosome, since the *E. coli* is conjugated to the pH-sensitive dye pHrodo green. After incubating the *E. coli* with the neutrophils, free *E. coli* are removed by pelleting and thoroughly washing the neutrophils. However, some *E. coli* remain bound to the surface of the neutrophils (depicted in the second neutrophil of Fig 1C), but they will fluoresce minimally in subsequent flow cytometry unless they are internalized into an acidic compartment. Upon addition of serum-opsonized NGs, these surface-bound *E. coli* become phagocytosed as bystanders when the NGs are phagocytosed (third neutrophil of Fig 1C). This leads to the *E. coli* particles accumulating in the low-pH phagosome, where they fluoresce during flow cytometry. Thus, the use of *E. coli* that fluoresce only in the phagosome allowed detection of a 2PAF > 1.

Having made the finding that nanogels can augment phagocytosis of bacteria, we tested whether this was a phenomenon unique to NGs. We performed experiments with NGs replaced as a “second particle” by a second *E. coli* particle, checking whether the second bacterial particles could enhance uptake of bacteria that were delivered during a first incubation. The experimental protocol was the same as Figure 1A, except that particle #1 was pHrodo green *E. coli,* and particle #2 was pHrodo red *E. coli* (Fig 2A). Here, the relevant conditions to compare are Figure 2C’s first vs. third green bars. This shows that phagocytosis of EC (non-serum exposed *E. coli*) is augmented by subsequent delivery of Serum EC (serum-opsonized *E. coli*). Indeed, the 2PAF = 1.65, or a 65% increase, (Fig 2C, inset, blue bar) is very similar to the 2PAF = 1.55 achieved with nanogels as particle # 2 (Fig 1C, inset, blue bar). Once again, the 2PAF is > 1 only when the first-delivered particle (EC) had not been opsonized. Thus, the NGs’ ability to augment uptake of bacteria is not unique to NGs, but is recapitulated in the neutrophil response to bacterial pathogens given in sequence. Our prior work shows that NGs, like bacteria, have tropism for neutrophils due to their rapid opsonization by complement (Myerson *et al.*, 2022). The parallel 2PAF effects for NGs and *E. coli* coincide with these similar uptake mechanisms.

**Figure 2.**
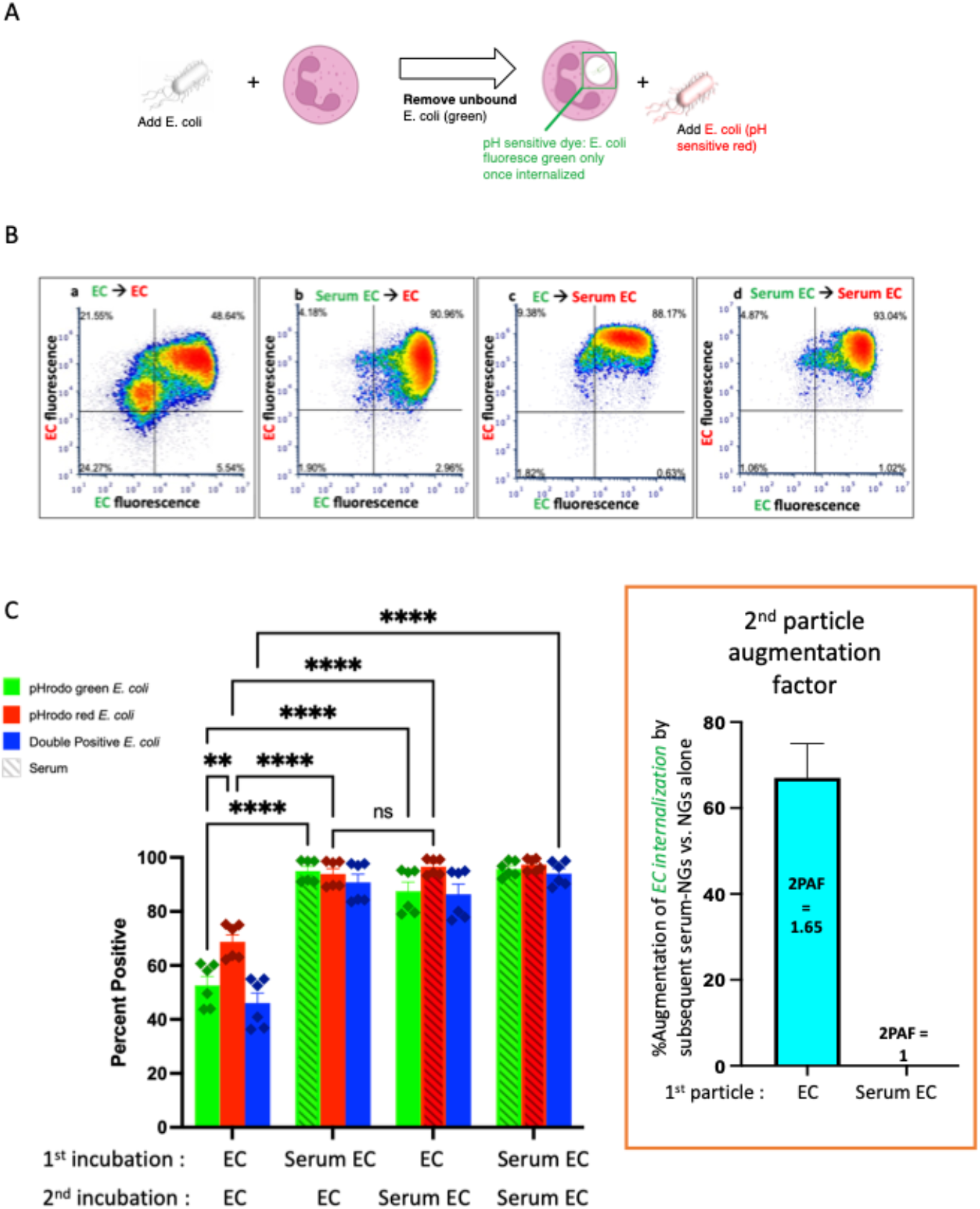
Similarly to serum-opsonized nanogels, serum-opsonized *E. coli* augment internalization of non-opsonized *E. coli.* A) Experimental scheme, similar to Figure 1A, but the second incubation was with *E. coli.* Note the first incubation was green E.coli, while the second was red *E. coli.* This experiment tests if, similar to serum-opsonized nanoqels, *E. coli* can augment phagocytosis of previously delivered *E. coli.* B) Representative dot-plots of the neutrophils, using the same notation as Fig 1. C) Quantification of flow cytometry experiments (n=6 biological replicates). Analogous to Fig 1, the most important comparison is the first vs third green bars, which both measure the % of neutrophils that phagocytosed non-serum-opsonized green *E. coli* (EC) which were present during the first incubation. When the second incubation was also with EC, only 50% of neutrophils were positive for the green EC presented during the first incubation. However, when the second incubation was with Serum EC, 85% of neutrophils were positive for the green EC presented during the first incubation. Thus, when the first particle is EC, the “2nd particle augmentation factor (2PAF)” was ~1.65 (inset, red). While Serum EC augment uptake of (non-opsonized) EC by 65%, Serum EC do not significantly augment phagocytosis of Serum EC that were present in the first incubation (2PAF = 1.0, blue bar in inset). Also notable is that Serum EC delivered as the first particle augments the uptake of EC delivered second (second red bar vs first red bar), showing the 2PAF effect does not depend on other order in which particles are delivered.

Having established that both serum-opsonized NGs and serum-opsonized bacteria can augment phagocytosis of non-opsonized bacteria, we questioned whether NG uptake could be similarly enhanced. We used the same protocol as above, except with particle #1 as NGs (green), and particle #2 as a separate sample of NGs (red) (Fig 3A). Comparing the first and third green bars in Fig 3C, we see that serum-opsonized NGs are *minimally* able to augment phagocytosis of *previously* delivered NGs. Thus, 2PAF = 1.23. These results may be attributed to difference in fluorophores of the *E. coli* bioparticles and NGs, as NGs fluoresce equally well on the surface of neutrophils (pH 7) and in phagosomes (pH 4-5). More likely though, the low levels of first incubation NG fluorescence observed in the first and third green bars suggest that neutrophils do not retain nanogels on their surface after washing, which would thus prevent augmentation by nanogels delivered during the second incubation. Figure 3C also shows that non-opsonized particle #2 NG uptake (the first red bar) is higher than non-opsonized particle #1 NG uptake (the first green bar). This implies that exposure of neutrophils to a first particle (even one that is mostly washed away) increases the phagocytic efficiency of the neutrophils for nanoparticles that are delivered later. This finding is consistent with previous studies showing neutrophils are known to change their activity state after phagocytosis(Bazzoni *et al.*, 1991). This finding suggests that the order and timing of particle delivery matter significantly.

**Figure 3.**
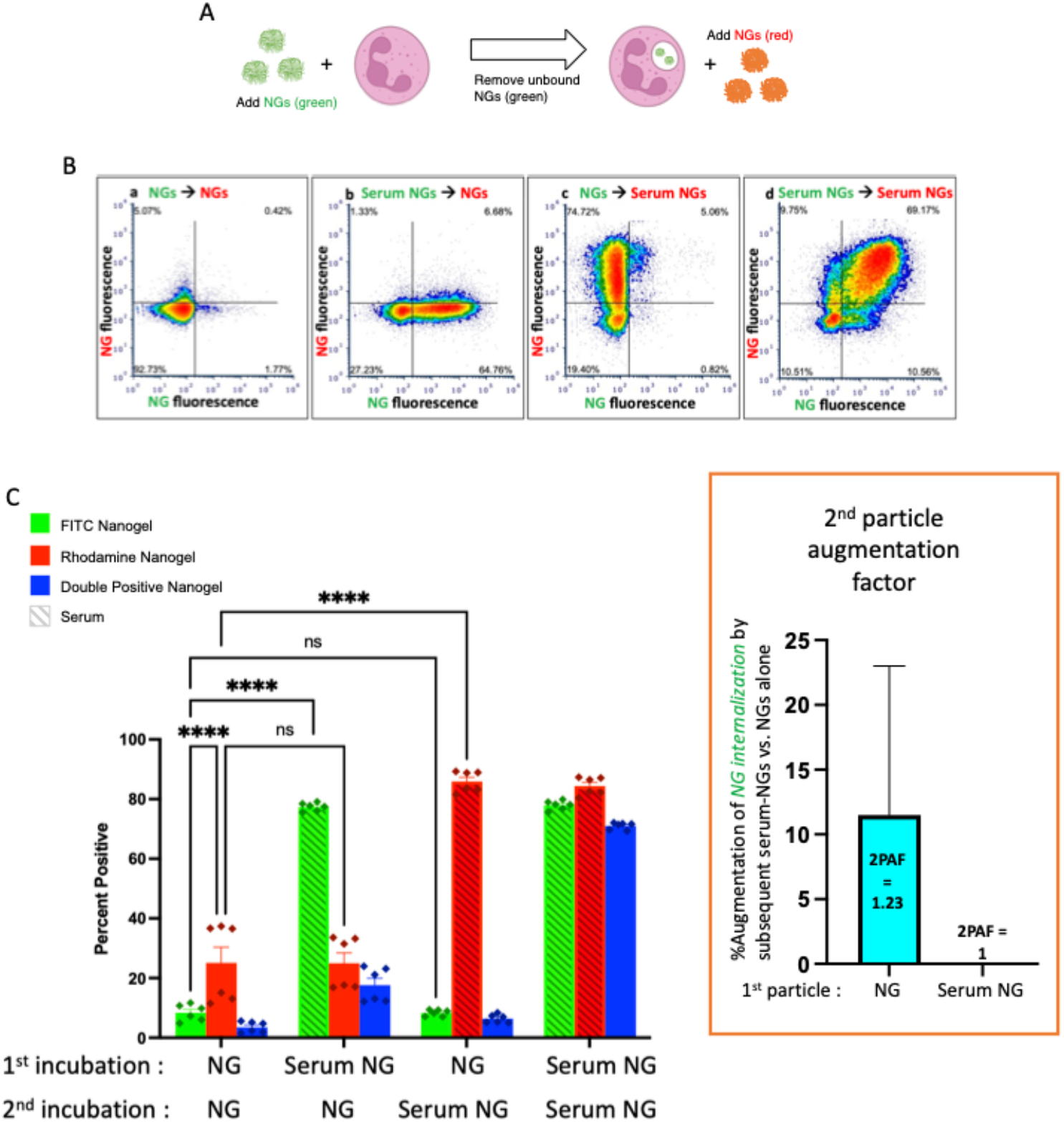
Serum-opsonized nanogels do not augment neutrophil association with previously delivered nanogels. A) Similar to the paradigm described in Figure 1, neutrophils were exposed to green nanogels, followed by washing to remove unbound neutrophils, then exposed to red nanogels, washed, and then the neutrophils were subjected to flow cytometry. B) Representative flow cytometry dot-plots. C) Quantification of n=6 biological replicates. The most notable comparison is the first and third green bars, which indicate that serum-opsonized nanogels (“Serum NGs”) do not augment the fraction of neutrophils that are positive for binding to non-serum-exposed nanogels (“NGs”). Notably, unlike the pH-sensitive *E. coli* used in Figs 1 & 2, nanogels fluoresce both when bound to the neutrophil surface and when in the phagosome. Thus, it is possible that some NGs are bound to the neutrophil surface and then internalized after exposure to Serum NGs, but this assay cannot detect such internalization events. Another major result of this set of conditions is that Serum NGs have uniformly high uptake into neutrophils regardless of whether the neutrophils were first exposed to other nanogels (NGs or Serum NGs), suggesting the neutrophils do not saturate their uptake of Serum NGs in this dynamic range (they do not “get full”).

Finally, we administered *E. coli* as particle #2 after NGs as particle #1 (Figure 4A). As in Figure 3, when NGs are delivered first, their uptake is minimally augmented by a second particle with 2PAF = 1.17, even when the second particle is highly stimulatory serum-opsonized *E. coli* (Fig 4C, first vs third green bars).

**Figure 4.**
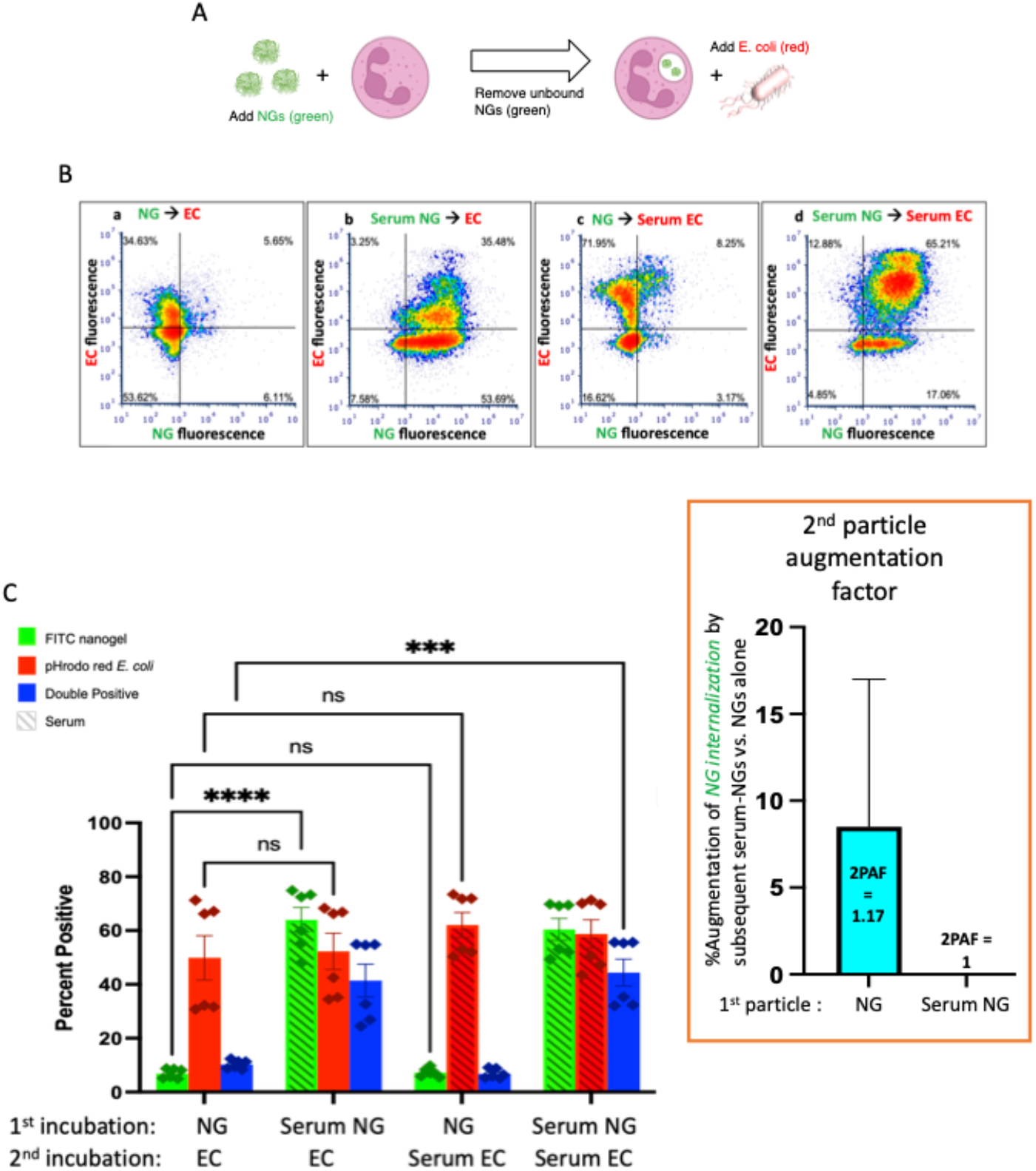
Serum-opsonized *E. coli* are also unable to augment neutrophil association with previously delivered nanogels. A) Here the first particle is nanogels, while the second particle is *E. coli.* Thus, this tests if EC can augment neutrophil association with nanogels. B) Representative flow cytometry dot-plots. C) Quantification of n=6 biological replicate. The most notable comparison again is the first and third green bars, which indicate that serum-opsonized *E. coli* (“Serum ECs”) do not augment the fraction of neutrophils that are positive for binding to non-serum-exposed nanogels (“NGs”). A second important result is that nanogels do *not* decrease neutrophil phagocytosis of *E. coli* (all red bars are equal). This is clinically relevant because it means that if a neutrophil phagocytoses a nanogel and then later encounters a bacterium, the nanogel will not reduce the likelihood of uptake of the bacterium. Thus, the nanogels do not induce immunosuppression by this metric.

Figure 4 also shows that delivery of NGs *before E. coli* does not impair neutrophils’ uptake of *E. coli:* levels of *E. coli* uptake were identical for all conditions tested in Fig 4C. This is an important result in the path to clinical translation of neutrophil-tropic nanoparticles, as such nanoparticles might compromise therapy if they inhibited subsequent phagocytosis of bacteria.

Taken together, the data presented here strongly indicate that serum-opsonized nanogels and *E. coli* augment phagocytosis of non-opsonized bacteria. We sought to confirm and extend these flow cytometry results with a complementary approach. Therefore, we performed a similar protocol of exposing neutrophils to nanogels and bacteria, but this time we analyzed the cells using microscopy. Not only could this serve as a confirmation of the flow cytometry results, but imaging data additionally can determine if the particles #1 and #2 co-localize with each other within the cell, which gives further insight into the 2PAF phenomenon.

We began this line of experiments by performing the same sequential delivery protocol used in flow cytometry experiments, but instead of flow cytometry, cells were fixed in suspension with paraformaldehyde and adhered to glass for microscopy imaging. Figures 5 A-C provide representative microscopy data. Fig 5A shows data where both particle #1 and particle #2 are *E. coli.* We found the same result as in flow cytometry: Serum EC delivered as particle #2 augmented neutrophil phagocytosis of (non-opsonized) EC delivered as particle #1 (compare green & yellow signals in Figs 5Aii vs 5Ai). Yellow signal in Fig 5A indicates intracellular overlap between green (particle #1) and red (particle #2) *E. coli* inside of neutrophils, consistent with a large fraction of particle #1 being phagocytosed at the same time and into the same compartment as particle #2. Figure 5B shows data where particle #1 is NGs (green) and particle #2 is *E. coli* (red). In this data, we observe minimal colocalization between the two particles, consistent with flow cytometry data in Figure 4 showing no 2PAF effect when NGs are given before *E. coli.* Figure 5C shows data where particle #1 is *E. coli* (green) and particle #2 is NGs (red). The condition for which flow cytometry yielded 2PAF = 1.55 (EC - Serum NG; [Fig 5Cii]), yielded imaging data with a high degree of intracellular overlap between *E. coli* (particle #1) and NGs (particle #2). These imaging data qualitatively support a key conclusion from our quantitative flow cytometry data: opsonized NGs or *E. coli* not only augment uptake of previously delivered, non-opsonized bacteria, but also show localization into similar intracellular compartments.

**Figure 5.**
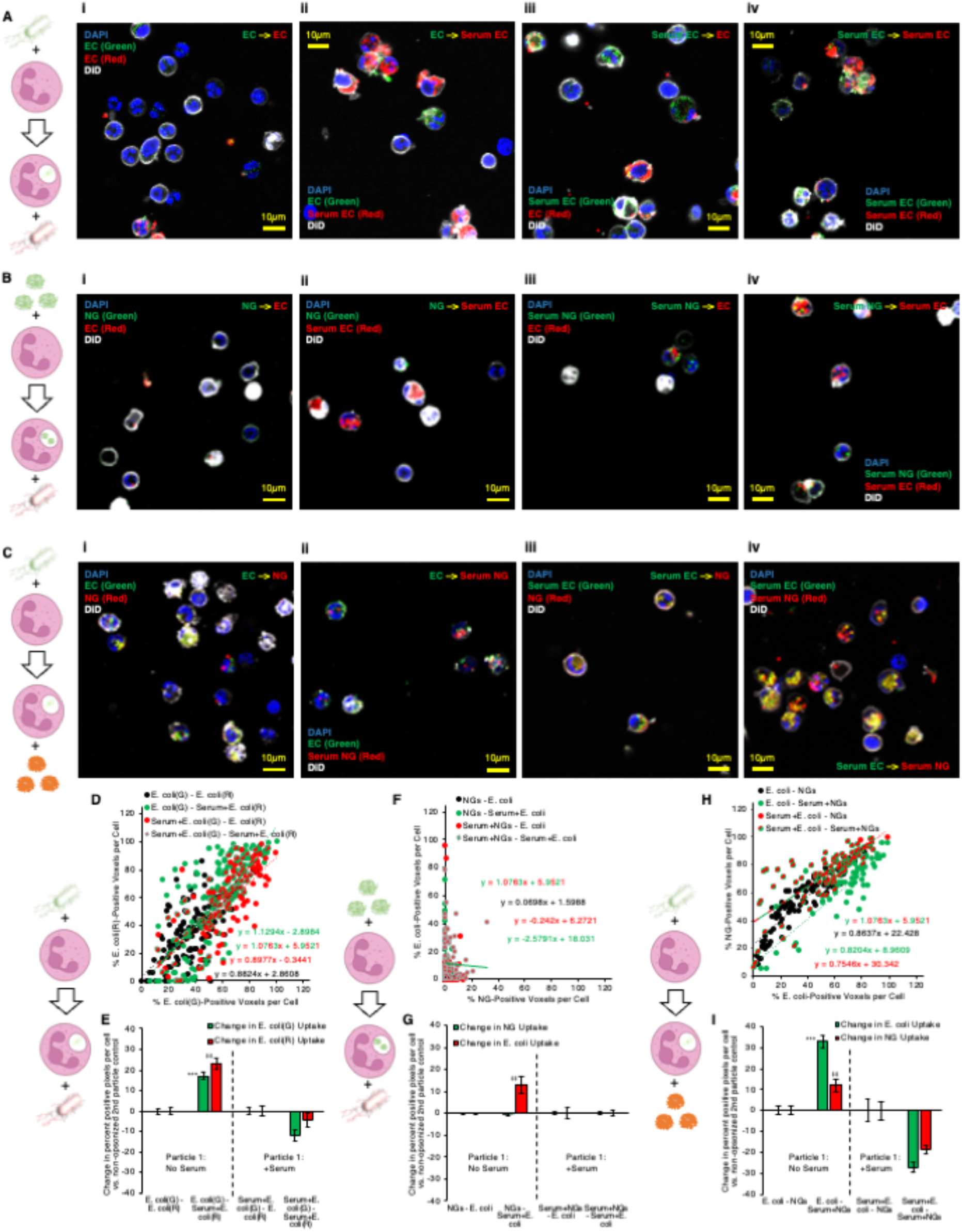
Microscopy confirms that serum-opsonized nanogels improve internalization of non-opsonized *E. coli,* and that they then share significant colocalization within neutrophils. A) Example images of neutrophils given two different labeled *E. coli* doses in sequence, for different serum pretreatment conditions applied to the *E. coli.* B) Example images of neutrophils given lysozyme-dextran nanogels (NGs) prior to *E. coli,* for different serum pretreatment conditions applied to the NGs and *E. coli.* C) Example images of neutrophils given *E. coli* prior to NGs, for different serum pretreatment conditions applied to the NGs and *E. coli.* Each point depicted in (D) indicates data for one cell, in imaging experiments as in (A). The horizontal coordinate indicates quantity of green *E. coli* signal and the vertical coordinate indicates quantity of red *E. coli* signal. Higher slope indicates that cells generally have more red *E. coli,* relative to quantity of green *E. coli.* All conditions showed strong positive correlation between red and green signal, indicating that cells taking up green *E. coli,* given first, were likely to take up red *E. coli,* given subsequently. E) 2PAF enhancement of first particle (green) *E. coli* by addition of serum to second particle (red) *E. coli.,* in imaging experiments as in (A). Bars 3-4 and 7-8 in (E) show the per-cell average difference in uptake of each color *E. coli* induced by serum treatment of second particle (red) *E. coli.* Green bars >0 indicate serum treatment of second particle (red) *E. coli* increases the average per cell uptake of first particle (green) *E. coli.* F-G) Data is as presented in (D-E), but for imaging experiments as in (B), wherein NGs were given to neutrophils before *E. coli.* There was poor correlation between per-cell NG signal and *E. coli* signal, for all serum pretreatment conditions, indicating that neutrophil uptake of NGs does not predict subsequent uptake of *E. coli.* G) indicates no 2PAF effect exerted by second particle *E. coli* on NGs. H-I) Data is as presented in (D-E) and (F-G), but for imaging experiments as in (C), wherein *E. coli* was given to neutrophils before NGs. In (H), there was strong correlation between NG and *E. coli* uptake in any given cell, when *E. coli* was taken up before neutrophil exposure to NGs. In (I), bar 3 indicates that opsonized second particle NGs enhance uptake of non-opsonized first particle *E. coli.*

To quantify the data from these microscopy experiments, images of each particle were thresholded via Renyi entropic filtering to identify the portions of each image that were positive for either particle #1 or particle #2. Guided by DiO membrane staining, we identified regions of interest encompassing each cell in each obtained image. Overlaying the thresholded images on the regions of interest corresponding to individual cells allowed us to determine the fraction of pixels in each cell that contained either particle #1 or particle #2. In panels 5D, 5F, and 5H, each point in the presented scatter plots represents the results of this analysis for one neutrophil. The y-axis value for each point represents the percentage of pixels in the neutrophil containing particle #2 and the x-axis value represents the percentage of pixels in the neutrophil containing particle #1. A line was fitted to the data for each condition. Lines with slope = 1 indicate that each neutrophil took up equal quantities of particle #1 and particle #2. Lines with slope > 1 indicate a tendency to take up more of particle #2 than particle #1. Lines with slope < 1 indicate a tendency to take up more of particle #1 than particle #2. We therefore used percentage of positive pixels as a metric for levels of particle #1 and particle #2 uptake in each neutrophil. To derive 2PAF values from the imaging data, we subtracted from our uptake values for each cell the average uptake values for the 2PAF baseline conditions (conditions where particle #2 is not opsonized). We therefore determined our 2PAF value in imaging experiments as the percent *increase* in either particle #1 or particle #2 uptake for each imaged cell vs. the expected level of uptake when particle #2 is not opsonized. These findings are depicted in panels 5E, 5G, and 5I.

Panels 5D and 5E depict quantitative analysis of imaging data where particles #1 and #2 were both *E. coli.* All serum treatment conditions in these experiments showed strong positive correlation between signal from particles #1 and #2, indicating that cells taking up green *E. coli,* given first, were very likely to take up red *E. coli,* delivered second. This confirms this assay behaves as expected: cells do not saturate their *E. coli* uptake in this dosing regimen; cells with more uptake of one *E. coli* have more uptake of the second; green and red fluorophores perform equally. As with flow cytometry data in Figure 2C, data in Figure 5E show a clear 2PAF effect exerted by particle #2 *E. coli* on particle #1 *E. coli.* When particle #1 *E. coli* is not opsonized, uptake of particle #1 increases by ~20% if particle #2 *E. coli* is opsonized, compared to data where particle #2 *E. coli* is not opsonized.

Panels 5F and 5G depict analysis of imaging data where particle #1 is NGs and particle #2 is *E. coli.* R-squared values were less than 0.1 for all lines in panel 5F except that for Serum+NGs – Serum+*E. coli,* indicating poor linear correlation between particle #1 and particle #2 uptake for the conditions in this data. This indicates that, when a given neutrophil takes up NGs as particle #1, improved uptake of *E. coli* as particle #2 cannot be predicted for that same cell. Similarly, analysis of imaging-based 2PAF also indicates no 2PAF effect exerted by particle #2 *E. coli* on particle #1 NGs. All imaging-based 2PAF calculations showed no change in particle #1 NG uptake induced by opsonized particle #2 *E. coli* vs. 2PAF baseline conditions with nonopsonized particle #2 *E. coli.* These data suggest that co-localization is compromised under these conditions, consistent with our understanding that multiple pathways can lead to phagocytosis, not all necessarily leading to accumulation in the identical intracellular compartment(Sahay, Alakhova and Kabanov, 2010).

Finally, panels 5H and 5I depict analysis of imaging data where particle #1 is *E. coli* and particle #2 is NGs. As with data in panel 5D, panel 5H shows strong positive correlation between particle #1 *E. coli* uptake and particle #2 NG uptake. For all serum treatment conditions, particle #2 NGs were more likely to be taken up in neutrophils that had already taken up particle #1 *E. coli.* This finding contrasts with the data in panel 5F, where, when NGs are given as particle #1, there was no positive correlation between NG uptake and *E. coli* uptake in any given cell. These findings match with analysis of the double-positive (*E. coli*-positive and NG-positive) cell counts in flow cytometry data in the blue bars in Figures 1C and 4C. Panel 5I shows imaging-based 2PAF values for conditions where *E. coli* is particle #1 and NGs are particle #2. Again, these image analysis findings agree with our flow cytometry data: When non-opsonized *E. coli* is particle #1, we observe a 33% increase in *E. coli* uptake per neutrophil when particle #2 NGs are serum-treated vs. when particle #2 NGs are not serum-treated. In imaging data, NGs as particle #2 exert a clear 2PAF effect enhancing uptake of *E. coli* as particle #1.

For conditions where we observed 2PAF effects (with *E. coli* as both particles #1 and #2 or with *E. coli* as particle #1 and NGs as particle #2), neutrophils were also examined with three-dimensional confocal imaging (Supplementary Figures 4 and 5). For our confocal imaging analysis, we computed quantities of NGs and *E. coli* in each cell as in Figure 5, but quantities of NGs and *E. coli* reflected fluorescent voxels, rather than pixels. Additionally, we quantified voxels that contained both particle #1 and particle #2, directly assessing three-dimensional colocalization of particles #1 and #2 in the confocal images.

For experiments where both particles #1 and #2 were *E. coli,* ~80% of *E. coli* particle #1 signal was spatially colocalized with *E. coli* particle #2 signal when *E. coli* particle #2 was opsonized (panels B-C, green bars 1 and 3 in panel C). Only ~50% of *E. coli* particle #1 signal was spatially colocalized with *E. coli* particle #2 signal when *E. coli* particle #2 was not opsonized (green bar 2 in Supplementary Figure 4, panel C). This finding fits well with a central part of our hypothesis as to the 2PAF mechanism: When there is a 2PAF effect, delayed phagocytosis of particle #1 is driven by coincident uptake of particle #2. In our confocal data, we find that particle #1 *E. coli* is mostly found colocalized with opsonized particle #2 *E. coli* inside neutrophils. This colocalization is diminished under conditions where 2PAF is diminished, when particle #2 *E. coli* is not opsonized. Under conditions with 2PAF effects, particle #2 colocalization with particle #1 (Supplementary Figure 4, panel A, red bars in panel C) was less than particle #1 colocalization with particle #2 (Supplementary Figure 4, panel A, green bars in panel C). This can be attributed to; a) opsonized particle #2 *E. coli* being phagocytosed independently of particle #1 at a higher frequency than events where particle #1 was phagocytosed independently of particle #2; b) opsonized particle #2 *E. coli* being taken up in neutrophils to a greater degree than particle #1 *E. coli.* Panel D in Supplementary Figure 4 indeed indicates higher average uptake values for *E. coli* particle #2 vs. *E. coli* particle #1, especially in 2PAF-affected conditions.

Our prototypical 2PAF conditions, where *E. coli* were particle #1 and NGs were particle #2, were also examined in confocal imaging data. Here particle #1 *E. coli* colocalized with particle #2 NGs at ~50-60% frequency when particle #2 NGs were opsonized (Supplementary Figure 5, panels B-C, green bars 1 and 3 in panel C). In comparison, *E. coli* colocalized with NGs at only ~35% frequency under non-2PAF conditions, where particle #2 NGs were not opsonized (Supplementary Figure 5, panels B-C, green bar 2 in panel C). As with data for 2PAF-affected conditions with double *E. coli* treatments, *E. coli-NG* 2PAF-affected conditions featured augmented colocalization of particle #1 (*E. coli*) with particle #2 (NGs), implying phagocytosis of particle #1 *E. coli* and particle #2 NGs in the same subcellular locations in neutrophils. Particle #2 NG colocalization with particle #1 *E. coli* (Supplementary Figure 5, panel A, red bars in panel C) was greater than particle #1 *E. coli* localization with NGs (Supplementary Figure 5, panel B, green bars in panel C). This may reflect generally higher uptake of *E. coli* vs. NGs (Supplementary Figure 5, panel D), where *E. coli* likely has a higher chance of being taken up in neutrophils independently of NGs, compared to lower quantities of NG uptake meaning NGs have a lower chance of being taken up independently of *E. coli.* Colocalization of particle #2 NGs with particle #1 *E. coli* bodes well for proposed NG drug delivery to neutrophils designed to help augment neutrophil killing of bacteria: When NGs are given after neutrophils have been exposed to bacteria, the NGs end up in the same places as bacteria inside neutrophils.

This study has several limitations. First, the data presented here are all derived from *in vitro* studies. Neutrophil isolation may modify neutrophil function, thus imposing constraints on interpretation. Of note, however, our previous study suggested a good correlation between *in vitro* and *in vivo* observations of NAP uptake in neutrophils (Myerson *et al.*, 2022). Similarly, the *in vivo* environment may contain neutrophil stimuli not seen *in vitro.* Opsonization with normal mouse serum also may not include all of the opsonins that might be present during an inflammatory process, although we have not detected any significantly different effect using acute phase sera (data not shown). Finally, neutrophils are significantly heterogeneous both within and between donors. Accordingly, use of multiple donors reduces this effect, and flow cytometry permits observation of potential heterogeneity, thus mitigating this concern.

## 3 Materials and Methods

### Lysozyme-Dextran Nanogel Synthesis

Lysozyme-dextran nanogels (LDNGs) were synthesized as previously described(Li *et al.*, 2008; Coll Ferrer *et al.*, 2014; Myerson *et al.*, 2018, 2022). Rhodamine-dextran or FITC-dextran (Sigma) and lysozyme from hen egg white (Sigma) were dissolved in deionized and filtered water at a 1:1 or 2:1 mol:mol ratio. Then pH was adjusted to 7.1 and solution was lyophilized. For Maillard reaction, the lyophilized product was heated for 18 hours at 60°C, with 80% humidity maintained via saturated KBr solution in the heating vessel. Dextran-lysozyme conjugates were dissolved in deionized and filtered water to a concentration of 5 mg/mL. Solutions were stirred at 80°C for 30 minutes. Diameter of LDNGs was evaluated with dynamic light scattering (DLS, Malvern) after heat gelation. Particle suspensions were stored at 4°C.

### Murine serum

Blood was obtained through the inferior vena cava as previously described(Mei *et al.*, 2010; Parasuraman, Raveendran and Kesavan, 2010) from B6 wild type mice and pooled. Blood was allowed to clot for 30 minutes at room temperature, and serum was separated by centrifugation of 1500 rpm x 10 minutes at 4°C. Complement proteins were depleted from serum via Cobra Venom Factor as previously described(Haihua *et al.*, 2018).

### Murine neutrophils

Femurs were harvested from B6 wild type mice, and bone marrow was collected and pooled. Neutrophils were isolated from bone marrow via negative selection (Stemcell EasySep™ Mouse Neutrophil Enrichment Kit cat #19762). Neutrophils were suspended in DMEM media at a concentration of 2 × 10^6^ neutrophils/mL. Isolated neutrophils were 95% viable by trypan blue, and 80% pure using Ly6g staining and flow cytometry (See Supplementary Figure 1).

### Nanogel preparation

FITC-labelled lysozyme-dextran NGs were synthesized as above. Stock NGs were brought to a concentration of 5 × 10^11^ particles/mL. To serum-treat prior to neutrophil incubation, NGs were incubated in 50% serum in DMEM for 1 h at 37°C.

### *E. coli* BioParticles

For bacterial particles, both pHrodo™ red and green *E. coli* BioParticles™ conjugates (Invitrogen, Thermo Fisher cat #P35361 and #P35366) were used. BioParticles were brought to a concentration of 2mg/mL in PBS. To serum-treat *E. coli* BioParticles prior to neutrophil incubation, BioParticles were incubated in equal volume serum for 1 h at 37°C.

### Prototypical Experiment

#### Sequential particle analysis

Neutrophils were isolated and prepared as above. 500uL of neutrophils were incubated with 20uL of either particle while rotating at 37°C. First particle incubation time was 15 minutes for NGs and 1 hour for *E. coli* BioParticles, with exception of the conditions with two color *E. coli* BioParticles. In those conditions the first *E. coli* BioParticles incubation was done at 15 minutes to recapitulate the initial NG incubation. Samples were then washed and pelleted 300g x 6 minutes to remove unbound particles. The neutrophils were then resuspended in 500uL DMEM media. The second particle was then incubated with the neutrophils while rotation at 37°C. Second particle incubation time was 15 minutes for NGs and 1 hour for *E. coli* BioParticles, with exception of the conditions with two color NGs. In those conditions the second NG incubation was incubated for 60 minutes to recapitulate the second exposure *E. coli* incubation. Samples were then washed and pelleted again, then resuspended in FACS buffer. For flow cytometry samples, they were stained with anti-Ly6G antibody prior to analysis. For microscopy samples, they were suspended in solution with 2% paraformaldehyde for 30 minutes at room temperature, then pelleted and resuspended at a concentration of 1 x 10^6^/mL.

#### Sample preparation and Confocal microscopy

90μL of paraformaldehyde fixed cells was incubated with 5 x 10^-7^ M DAPI and 1:100 Vybrant™ DiD Cell-Labeling Solution (Invitrogen) at 37C for 20 minutes. Samples were centrifuged at 2000g x 1 minute. They were washed with 1mL PBS and sprun at 2000RPM x 3 minutes twice. Finally, they were resuspended in 80μL PBS then dropped on cavity slides (Eisco), covered with coverslips (FisherScientific), and analyzed with Leica TCS SP8 Laser confocal microscope. Visualization of neutrophils was performed with water immersion objective HC PL APO CS2 40x/1.10. Images were obtained in sequential scanning mode using Diode 405, OPSL 488, OPSL 552 and Diode 638 lasers. We used scan speed 200 Hz and pixel size 0.223 μm for flat images. Z-stacks were obtained with scan speed 600 Hz, XY pixel size 0.223 μm and voxel size 0.424 μm. Images and Z-stacks were processed with LASX (Leica microsystems). Images and Z-stacks were converted to TIFF images for analysis in ImageJ (FIJI distribution 2.1.0/1.53q). Processing and analysis, including image thresholding, fluorescence colocalization, and per-cell analysis of fluorescence signals, employed custom ImageJ macros, code for which is provided in full in the supplement. For per-cell analyses, regions of interest were drawn manually around each imaged cell, using images of DiD membrane stain to define the edges of individual cells.

#### Animal study protocols

All animal studies were carried out in strict accordance with Guide for the Care and Use of Laboratory Animals as adopted by National Institutes of Health and approved by Children’s Hospital of Philadelphia Institutional Animal Care and Use Committee. All animal experiments used male B6 mice, 6–8 weeks old, purchased from Jackson Laboratories. Mice were maintained at 20–25 °C, 50% ±20% humidity, and on a 12/12 h dark/light cycle with food and water ad libitum.

#### Statistical analysis

Error bars indicate the standard error of the mean throughout. Significance tests are described in captions. Statistical power was determined for statements of statistical significance and tabulated in the supplementary materials.

## 4 Conclusion

The data presented here are consistent with a model of neutrophil phagocytosis in which a non-opsonized bacteria, that is poorly phagocytosed, is nonetheless still available in a compartment (likely the surface plasma membrane) from which it can be subsequently taken up in response to a more phagocytic-stimulatory particle, whether it be an opsonized bacteria or an opsonized nanogel. Similarly, initial exposure to a bacterial-like particle further enhances subsequent uptake of nanogels, even non-opsonized ones. Thus, the nanogel might be used in its opsonized form, even without incorporated drugs, to enhance uptake of poorly-opsonized bacteria. Furthermore, under these circumstances, the bacteria and nanoparticles are found in similar intracellular compartments, suggesting that delivery to specific compartments of the phagocytosing neutrophil might be possible.

## Acknowledgements

We are grateful for the support by the grants from the National Institutes of Health F32-HL-151026 (KMR), RO1-HL-151467 and RO1-HL-158737 (VPK), R01-HL-157189 (VRM/GSW), K08-HL-138269, R01-HL-153510 and R01-HL-160694 (JSB), and UH3-TR-002198 (GSW), as well as, the Job Research Foundation Grant (VPK).

