## Supplemental figures for "Nanoparticle-induced augmentation of neutrophils’ phagocytosis of bacteria"

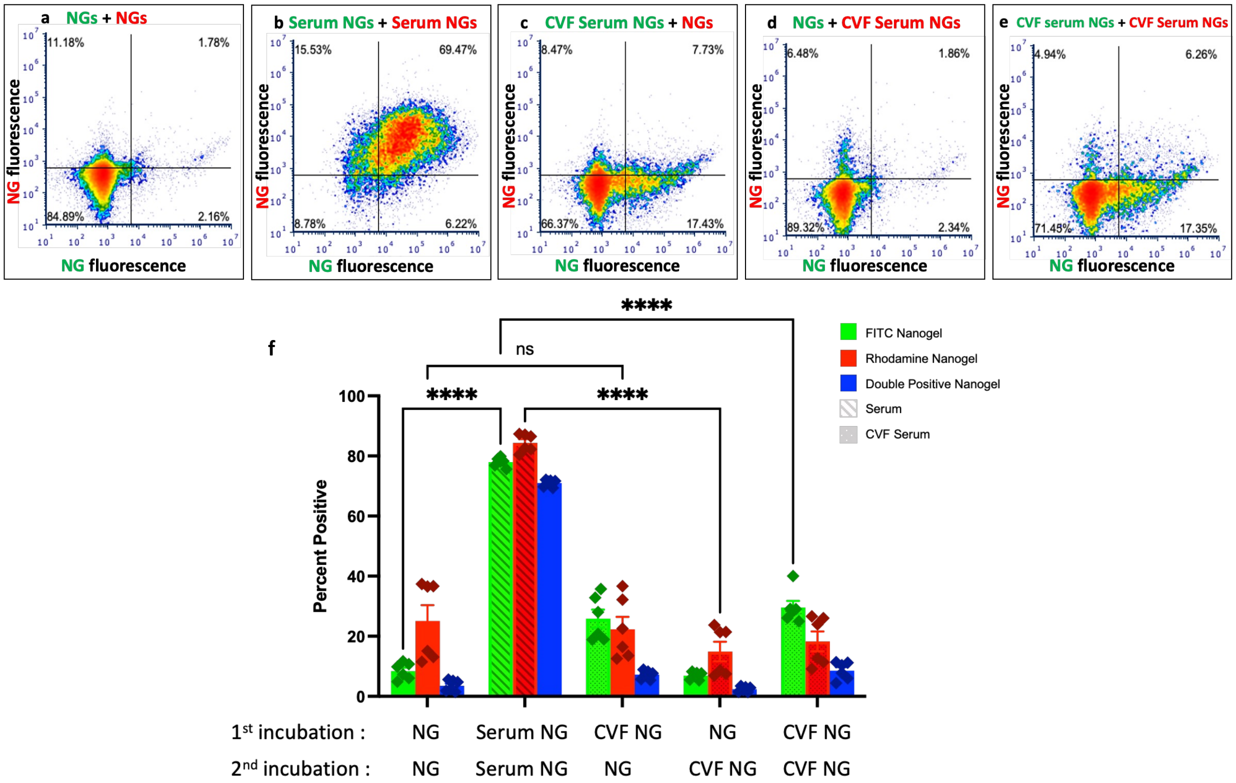


**Supplementary Figure 1.** *Cobra venom factor (CVF) treatment ameliorates the increased NG uptake seen as a serum effect.* Panels a-e show representative flow cytometry plots, and panel f is a summative representation of data. (a) After sequential incubation with unopsonized FITC (green) NGs and then rhodamine (red) NGs, Neutrophils show a low positivity for each particle. As compared with (b) where NGs were incubated with wild type serum for 1 hour prior to neutrophil incubation. There is significant shift in the percent of positive neutrophils. There is additionally a shift in fluorescence with cells in (a) concentrated around $10^3$ fluorescence but in (b) are seen up to $10^6$ fluorescence, a 1000-fold increase. For panels c-e, CVF was used. CVF is a complement-activating protein that depletes C3 in serum. Serum was exposed to CVF for 60 minutes and then spun to remove any denatured proteins prior to incubation with NGs. In (c) and (d), where one particle was incubated in CVF-treated serum, the flow plot does not completely recapitulate panel (a) to where they look unopsonized; however, there is a significant decrease in the serum effect seen previously. (e) additionally shows some increased cell positivity as compared to (a), but as compared to it's double serum exposed NGs analog in (b) there is amelioration of the serum effect. These findings are also appreciated in (f). When comparing the third and fifth green bar (CVF NGs) to the first two, there is a significant decrease in the percent of positive neutrophils without fully reaching the low level of the unopsonized (first green bar) NGs. (*p<0.05)

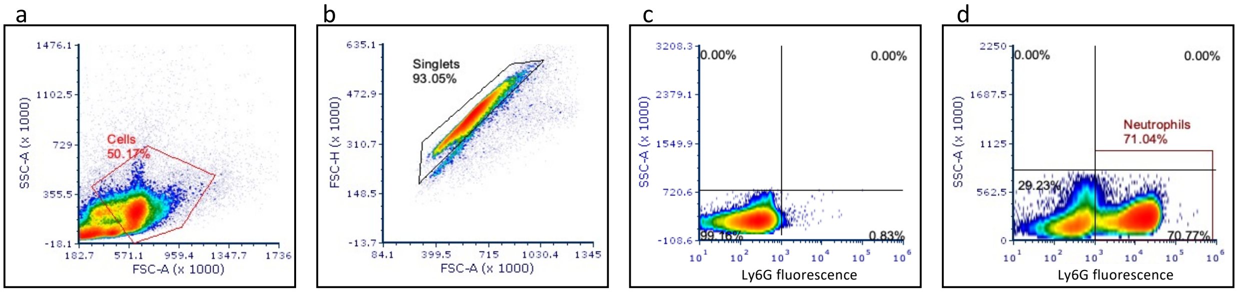


**Supplementary Figure 2.** *Flow cytometry cell gating strategy.* Following neutrophil enrichment by negative selection, an unstained condition and a PerCP-Cy5.5 anti-Ly6G stained single color condition were prepared for flow cytometry. (a) Unstained cells were used to gate out debris. (b) From the ”cell” gate of (a), the unstained singlet cells were then gated in for subsequent plots. (c) Quadrant gating for Ly6G positivity was determined on the cells within the “singlet” gate of an unstained condition. (d) All cells that stained for Ly6G were gated into the “neutrophil” gate and used as the gate for the analysis of NG and *E. coli* bioparticle uptake.

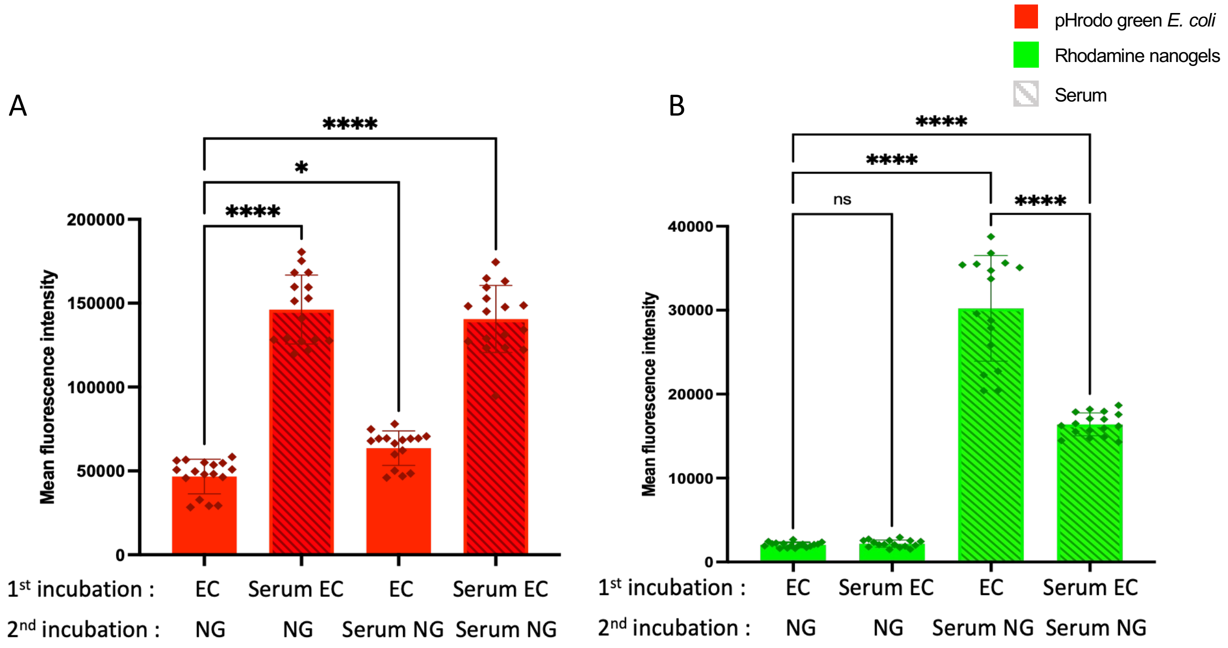


**Supplementary figure 3.** Mean fluorescence intensity data from the experiments in presented in Figure 1. A shows the MFI of particle #1, *E. coli*. When comparing the first and third bars, we again see the 2PAF effect where more pHrodo green fluorescence is detected when particle #2 is a serum-opsonized NG. B shows the MFI of particle #2, NGs.

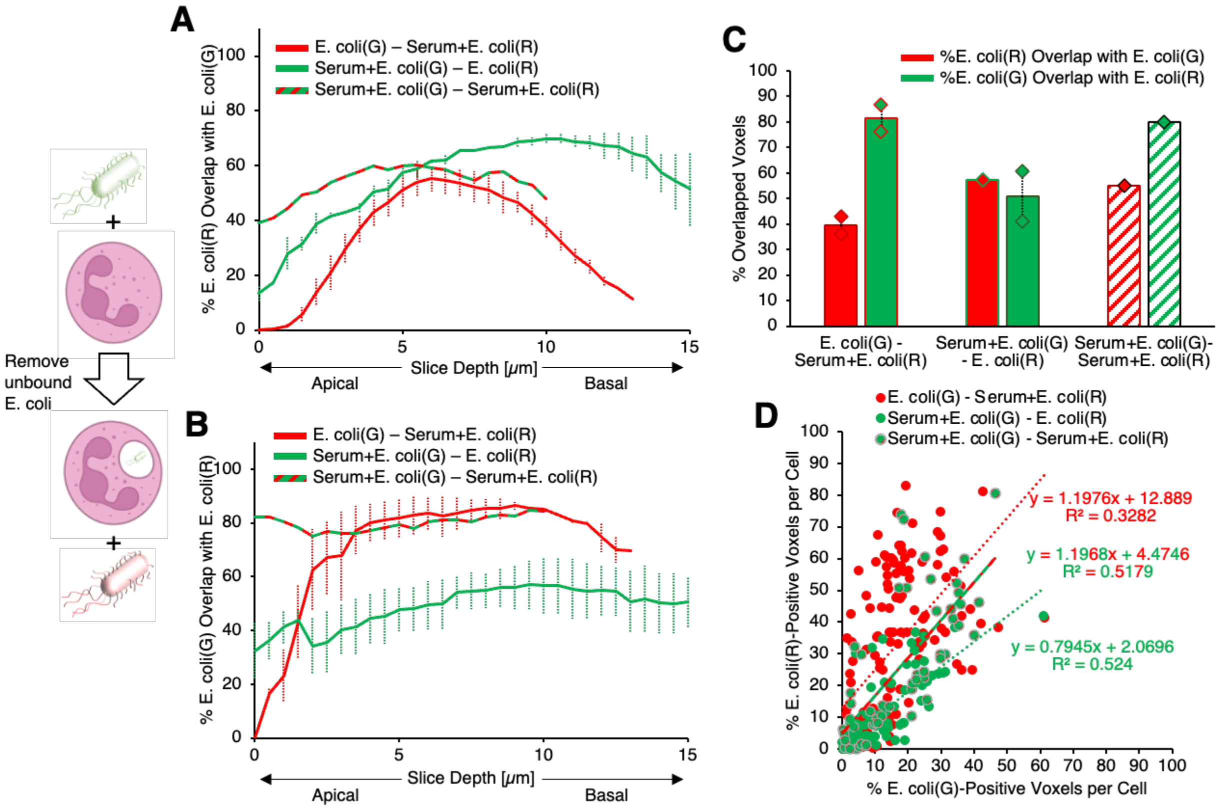


**Supplementary figure 4.** In experiments with red E. coli given to neutrophils after green E. coli, confocal imaging data indicated colocalization of red/second dose E. coli with green/first dose E. coli (a) or colocalization of green/first dose E. coli with red/second dose E. coli (b) as a function of depth in plated cells with different E. coli serum pretreatment conditions. Identical data is condensed over all depths in (c). (d) depicts, for each imaged cell, quantity of red/second dose E. coli vs. quantity of green/first dose E. coli under different serum pretreatment conditions, showing correlation between uptake of particle 1 E. coli and uptake of particle 2 E. coli for each imaged cell.

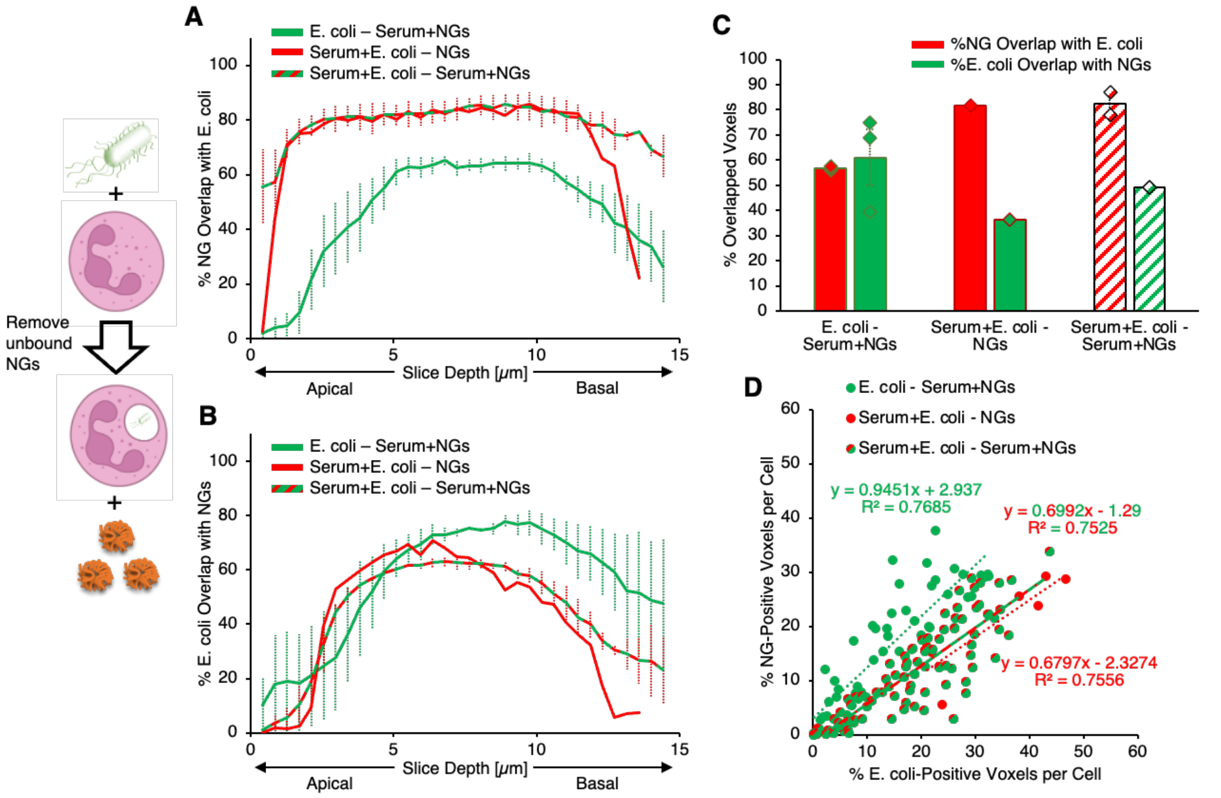


**Supplementary figure 5.** In experiments with red NGs given to neutrophils after green E. coli, confocal imaging data indicated colocalization of NGs with E. coli (a) or colocalization of E. coli with NGs (b) as a function of depth in plated cells with different E. coli and NG serum pretreatment conditions. Identical data is condensed over all depths in (c). (d) depicts, for each imaged cell, quantity of E. coli vs. quantity of NGs under different serum pretreatment conditions, showing correlation between uptake of particle 1 E. coli and uptake of particle 2 NGs for each imaged cell.
